# Defining region-specific masks for reliable depth-dependent analysis of fMRI data

**DOI:** 10.1101/557363

**Authors:** Kimberly B. Weldon, Philip C. Burton, Andrea N. Grant, Essa Yacoub, Cheryl A. Olman

## Abstract

In high-field fMRI research, anatomical reference information (e.g., gray matter (GM) segmentation, cortical depth delineation) is often defined in volumes acquired with pulse sequences subject to different distortions than those in functional volumes. In these cases, reliable interpretation of ultra-high resolution fMRI data depends on excellent cross-modal registration of functional volumes to reference anatomical volumes. In this paper, we describe a two-step approach to automating assessments of cross-modal registration quality for the purpose of guiding depth-dependent analysis. First, each functional/anatomical registration was scored by the ratio of the number of GM voxels in the functional data overlapping anatomical GM to the number of GM voxels in the functional data overlapping anatomical white matter (WM). This GM:WM overlap ratio provided an objective metric for determining whether an alignment algorithm had converged on a solution that would pass visual inspection. Second, surface-based maps indicating the consistency of overlap between functional and anatomical GM throughout the GM depth were generated for the entire region of cortex covered by the experiment. These maps served as a mask for the purpose of excluding regions where registration between functional and anatomical data was inadequate and thus unable to support depth-dependent analyses. We found, for both real and simulated data, that functional response profiles that were less biased toward superficial responses in regions where these metrics indicated satisfactory registration.

## Introduction

The improving technology of high-resolution functional magnetic resonance imaging (fMRI) enables examinations of sub-millimeter spatial maps of brain function. In particular, this sub-millimeter resolution allows investigations into the laminar specificity of the hemodynamic response in humans [1–9]. However, fMRI data are typically acquired with echo-planar imaging (EPI) readouts, which are particularly sensitive to B_0_ field inhomogeneities, resulting in distortions in the images themselves [10]. In addition to these distortions, EPI images have relatively poor contrast between gray matter (GM) and white matter (WM). To combat these problems, it is common for information about cortical region and depth to be derived from T_1_-weighted reference images [e.g. MP-RAGE, 11], which are not as susceptible to distortion and have improved contrast between GM, WM, and cerebral spinal fluid (CSF). As a result, quality cross-modal alignment of functional and anatomical volumes is of particular importance in the analysis of high-resolution fMRI data.

Efforts to perform GM/WM segmentation directly on distorted functional data [12] or on anatomical volumes that are acquired to be distortion-matched to fMRI data [13,14] are of course a superior alternative to cross-modal registration [see 15 for a recent discussion]. However, acquiring distortion-matched volumes with adequate contrast and coverage to support segmentation is not always an option, either because the necessary, specialized pulse sequences are not available on a scanner or because the time required to generate this extra set of images makes an experimental session too long. Therefore, improving cross-modal registration methods for the purpose of translating of segmentation information from one dataset to another remains valuable.

The present work remains agnostic about the best ways to perform GM/WM segmentation (e.g., with or without the aid of a T_2_-weighted image to refine the pial surface [16], whether higher resolution than 1 mm is required for accurate segmentations, and how cortical depth should be computed [17]). For simplicity, here we use the default FreeSurfer (http://surfer.nmr.mgh.harvard.edu/) segmentation on a T_1_-weighted MP-RAGE [11] acquired at 3 Tesla with 1 mm isotropic resolution. Anatomical segmentation is, of course, better with higher-resolution T_1_-weighted images supported by matched T_2_ acquisitions. However, the occasional error at the pial surface or failure to mark the depth of a sulcus provides illustrative examples and does not impact the overall results for the study at hand.

It is also important to note that the amount of distortion in the functional data varies widely between datasets, and there are many different methods for doing distortion compensation. Here, we use AFNI’s `3dQwarp` tool to apply distortion compensation using a blip-up/blip-down approach, which is one of several (e.g., FSL’s `topup` and `FUGUE`) suitable choices. It is safe to assume that, while all laboratories work to minimize distortion during data acquisition and preprocessing, all functional data will retain some residual distortions and blurring. The emphasis in this work is on quantifying the impact of these effects.

## Step 1: The GM:WM overlap ratio (GWOR) as a metric for overall registration quality

Assessing the quality of registration between functional and anatomical data has historically involved visual inspection by human observers. Visual inspection is inherently subjective and time-consuming, making it impractical for use in, for instance, whole brain analyses, on a large number of datasets in a given experiment, and especially in instances where the GM/WM image contrast in the functional data is so poor that segmentation on the functional dataset itself is not feasible. In this section, we present a pipeline for assessing the overall quality of a cross-modal registration that largely avoids the need for visual inspection.

First, we mark the GM in the functional volume. Then we calculate the number of functional GM voxels that overlap anatomical GM, as well as the number of functional GM voxels that overlap anatomical WM. The ratio of these two numbers (the GM:WM overlap ratio, GWOR) can be used as a score for the purpose of determining relative quality of the overall alignment.

In pursuit of this metric, we compare two different practical and readily available methods of marking the GM in the functional data. In the first approach, we perform image-based segmentation on the functional data to define GM, WM and CSF classes. In the second approach, we create a binary mask using the activation from a robust, independent functional localizer performed during the same scanning session and acquired with the same image acquisition parameters as the main functional task. This second, novel approach is based on the assumption, evident in our data, that neural activation is largely present in the GM as opposed to the WM [but see 18,19].

Using activation as a proxy for GM in the functional images raises concerns about bias stemming from strong responses in pial vessels and weak responses in deep GM. In this section, we also explore the confounds that might arise from unwanted signal modulation in large surface (pial) vessels and from the difficulty in detecting responses near the GM/WM boundary by applying these approaches on a simulated dataset.

### Step 1: Material and methods

#### Participants

After providing written informed consent, three healthy adult subjects (one male) participated in each of two experiments with different visual stimuli but identical data acquisition strategies (6 datasets total). The experimental protocol was approved by the University of Minnesota Institutional Review Board.

#### Data acquisition

Functional MRI data were collected at the University of Minnesota’s Center for Magnetic Resonance Research on a Siemens 7 Tesla scanner equipped with a SC-72 body gradient set (70 mT/m; 200 T/m/s) using GE EPI at 0.8 mm isotropic resolution (matrix size = 168 × 200; multiband = 2; TR = 2000 ms, TE = 22.6 ms, 56 coronal slices, R/L phase encode (PE) direction). Data were acquired with a parallel imaging acceleration factor of 3 (6/8 Partial Fourier; echo-spacing = 1.0 ms). We used a custom 4-ch transmit, 32-ch receive head coil. The relative phases of the coil were constant for all participants and achieved a B_1_ transmit solution [20] that provided adequate power throughout early visual regions.

In the first experiment, each participant passively viewed dichoptically-presented clips of an animated movie (12 scans, 12 s to the left eye /12 s to the right eye, 10.5 cycles per scan). In addition, there were four independent localizer scans. Each localizer scan was an adapted population receptive field (pRF) mapping scan [21] during which participants fixated on a central fixation point during the presentation of moving bars that featured highly-salient stimuli (e.g. faces, body parts, objects, or animals; 4 scans, 16 bar sweeps per scan at 8 orientations and 2 directions).

In the second experiment, participants viewed dichoptically-presented movie clips (4 scans, 12 s to the left eye /12 s to the right eye, 10.5 cycles per scan) interspersed with rest blocks as well as dichoptically-presented flickering checkboards (4 scans, 10.5 cycles per scan). The same four pRF scans from Experiment 1 were included in Experiment 2, but the movie-watching scans provided more robust responses and were used as the independent localizers for generating laminar profiles from the checkerboard scans.

During each experimental session, functional volumes with reversed phase-encoding polarity (30 TRs, L/R PE) were acquired in each experimental session as well as anatomical scans (“Within-Session T_1_”, 1 mm isotropic T_1_-weighted MP-RAGE) to guide registration. Cortical surface definition and segmentation were performed on anatomical volumes (“Reference T_1_”, 1 mm isotropic T_1_-weighted MP-RAGE) acquired in separate scanning sessions on a 3 Tesla system in the same facility. Due to reduced inhomogeneity in the B_1_ field, images acquired at 3T display less intensity variation than 7T data, which can result in improved segmentation [22].

#### Preprocessing

Functional runs from each dataset were motion-compensated using AFNI’s `3dvolreg` tool. Next, each dataset was distortion-compensated using AFNI’s `3dQwarp` tool (-`plusminus` flag) to generate a voxel displacement map from EPI images acquired with opposite phase encode directions. Distortions in the functional data due to known imaging gradient non-linearities were also compensated using the grad_unwarp toolbox provided at https://github.com/Washington-University/gradunwarp. WARP maps (from both gradient non-linearity correction and B_0_ distortion correction) were combined with motion-compensation matrices to create preprocessed data subjected to only one resampling step. Gradient non-linearity correction was also applied to the Reference T_1_ and the Within-Session T_1_.

We used AFNI’s `3dAllineate` command to register the Within-Session T_1_ to the participant’s Reference T_1_. This procedure generated a transformation matrix that was used to define an optimal starting point for the affine registration between the functional datasets and the Reference T_1_. GM/WM segmentation was performed on the Reference T_1_ using FreeSurfer (`recon-all` command, using all defaults).

#### Affine registration between functional and anatomical data

We used two different alignment tools and several different cost functions for a total of six different rigid-body, intensity-based registration algorithms to determine the rotation and translation that would provide an affine alignment between each participant’s functional dataset and their Reference T_1_. The goal of using several different registration algorithms was not to provide an exhaustive comparison of available algorithms, but to use a representative sampling that would produce a wide range of results. For registrations using AFNI’s `3dAllineate` tool, we used three different cost functions: correlation ratio (`crM`), normalized mutual information (`nmi`), and a cost function based on maximizing negative correlations between functional and anatomical data [lpc; 23]. For registrations using FSL’s `FLIRT`, we used two different cost functions: the default correlation ration (`corratio`, which uses positive or negative correlations between intensity values, and boundary based registration (`bbr`), which uses the intensity difference across the WM boundary in the reference image. All of the registrations relied on a binary *epi* mask as a weighting function, indicating which voxels in the functional data should be used when computing the cost function.

#### Marking functional GM using image-based segmentation

We created a marker of functional GM using imaged-based segmentation (GM_seg_). After an initial (and plausible, by visual inspection) registration between the functional volume and the Reference T_1_ was accomplished, the cortical parcellation provided by FreeSurfer was resampled to the functional space and provided to AFNI’s `3dSeg` as an initial segmentation (using the `-cset` flag). The output of this process is a ternary mask with CSF labeled as 1, GM as 2, and WM as 3. A sample EPI segmentation is shown in Fig. 1.

**Fig 1.**
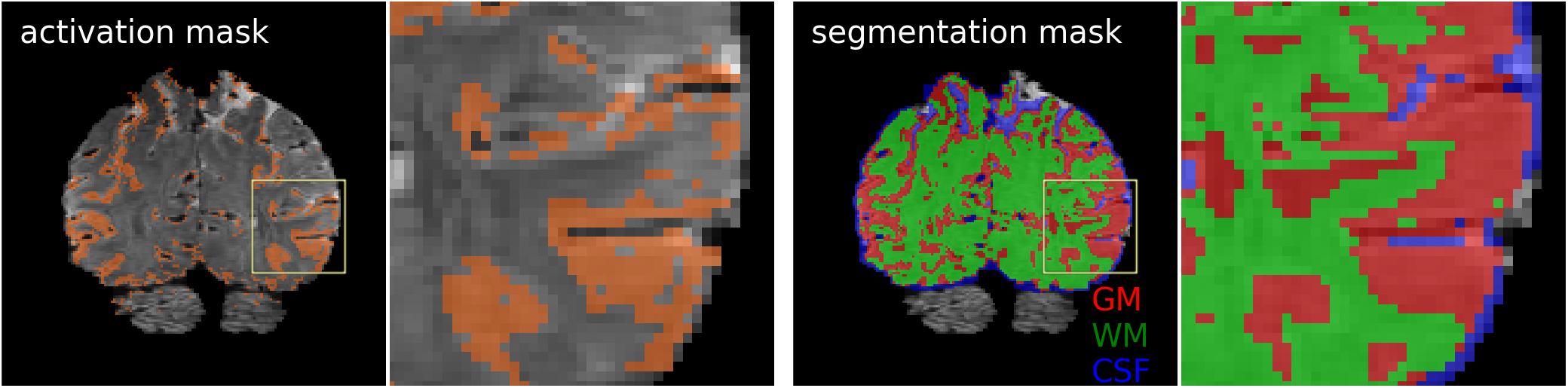
Two different methods of marking functional gray matter (GM). Left two panels: GM_act_, using activation from an independent localizer (grayscale underlay: EPI data; orange overlay: binary activation mask, showing all voxels correlated with stimulus presentation at p (uncorrected) < 10^−6^). Right two panels: GM_seg_, using image-based segmentation by AFNI’s `3dSeg` after initializing segmentation with GM, WM and CSF masks translated from anatomical to functional space (red: GM; green: WM; blue: CSF). Limitations to both approaches are detailed in the text.

#### Marking functional GM using activation from an independent localizer

In another approach to marking functional GM, we created binary activation masks using a standard general linear model (GLM) for pRF scans in Experiment 1, or a Fourier analysis for the block-design independent localizers in Experiment 2 (GM_act,_ Fig. 1). An individual voxel threshold of *p* < 10^−6^ (uncorrected) was used to create a binary activation mask. Pial vessel contributions to these masks were minimized by removing voxels in which the functional image signal-to-noise ratio (mean divided by standard deviation) was less than 10. Previous work [24] has verified that mean-normalized variance (the inverse of SNR) is a good metric for large vessels [for a similar approach, see 2].

#### Computation of GM:WM overlap ratio (GWOR) for two functional GM markers

Following these computations, each of the six functional datasets had six different registrations, for a total of 36 possible registrations to assess, either by visual inspection or by computing a GWOR score. A given GWOR score was computed by taking the ratio of (1) the number of voxels in the functional GM mask and (2) the number of voxels in the anatomical GM mask (as defined in the `wmparc` file from FreeSurfer and then translated to the functional space by the appropriate registration). Because we had two different GM markers (GM_seg_ and GM_act_), two different GWOR scores were available for each registration (GWOR_seg_ and GWOR_act_).

#### Visual inspection of registration quality

The use of multiple registration cost functions on six datasets resulted in 36 possible registrations. However, two of the alignment algorithm/cost function combinations (`3dAllineate/nmi` and `FLIRT/corratio`) reliably produced non-viable solutions (no overlap between functional and anatomical data) for our datasets; those registrations were discarded.

Five expert observers (including three of the authors) evaluated the quality of the remaining 24 registrations on a scale of 0-5. A score of 0 represented images for which registration appeared to fail completely (no overlap between functional and anatomical data), and a score of 5 indicated the appearance of a perfect registration. Images were presented in random order to the observers so as not to bias their responses toward a particular subject, experiment, or registration technique.

#### Simulation

Four simulated datasets were created to test the underlying assumptions of the analysis approach described above (Fig. 2). First, a simulated *T_2_*-weighted EPI* dataset was created (Fig. 2A). The GM and WM masks created by FreeSurfer (available in the `wmparc` file) were resampled into a volume with the same resolution and field of view as the real EPI data. GM (and all of cerebellum) was assigned a value of 500 and WM was assigned a value of 650. Then, a dilated version of the stripped brain was used to create a CSF mask; all voxels in the mask that were not GM, cerebellum or WM were assigned a value of 1000.

**Fig 2.**
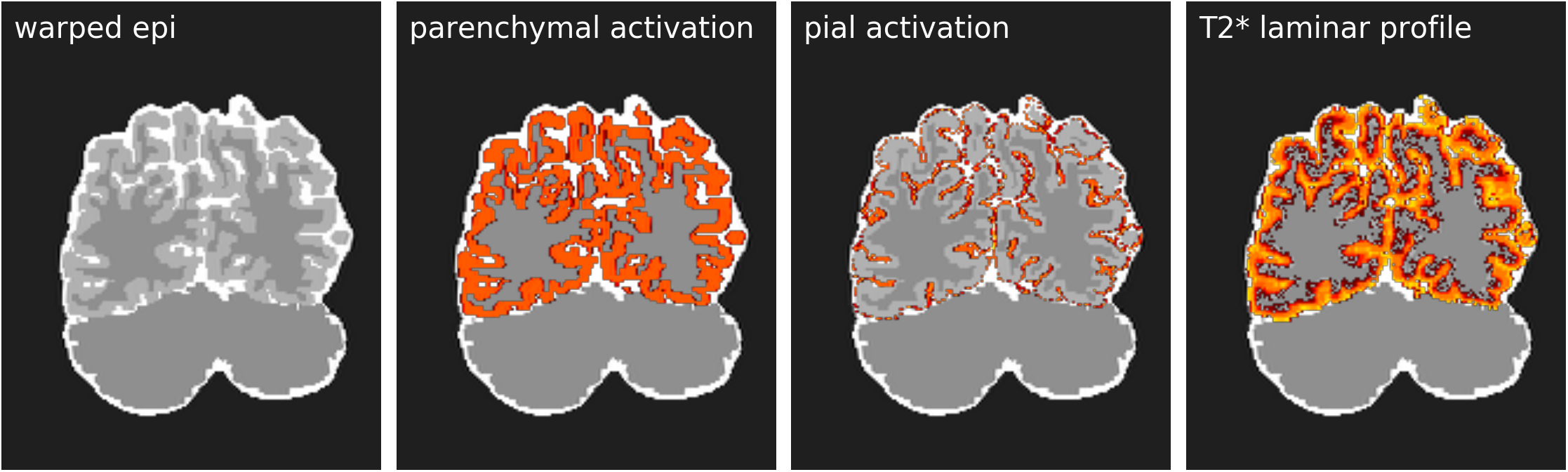
Simulated datasets. Three different T_2_*-weighted volumes (one for each of the three unique participants in this study; one slice of one simulation shown here) were simulated by resampling each reference anatomy to match the resolution and slice prescription of the real data. Simulated activation patterns were created using information from the GM segmentation and pial surface definition in the reference anatomy (see text for details). The simulated datasets were warped using the actual voxel displacement maps from the real experiments, and passed through the same analysis pipeline as the real data.

Next, three simulated activation profiles were created. A ground-truth GM mask was created by assigning a value of 1 to GM voxels and 0 elsewhere. This approach simulated a robust binary activation mask from an independent localizer limited to the parenchyma (*parenchymal activation*, Fig. 2B). A binary mask was created from the pial surface, simulating a worst-case scenario of activation only in pial vessels (*pial activation*, Fig. 2C).

Finally, a plausible T_2_*-weighted activation profile, constrained to the GM, was simulated by assigning 0.3% signal change to voxels near the WM boundary, 1.4% to voxels at the pial surface, and an essentially linear ramp in between the two boundaries, with a moderate elevation of responses in the middle of the cortical thickness. To simulate the problem of activation being harder to detect in deep GM where responses are lower, random noise (uniformly distributed between −0.25 and 0.25) was added to the data. The *T2* laminar profile* (Fig. 2D) was finalized by thresholding the data at 0.3 (thereby disproportionately affecting “deep” responses). Noise levels of 0, 0.25 (shown) and 0.5 were tested, and the only effect was a decrease in estimated response amplitude in the deepest parts of GM (as would be expected if a smaller proportion of the voxels were responsive).

All four simulated volumes (*T_2_*-weighted EPI, parenchymal activation, pial activation, and T2* laminar profile)* were warped using the voxel displacement masks measured during the real experiments. The simulated, *warped EPI* was registered to the Reference T_1_ using AFNI’s `3dAllineate` (`lpc` cost function; *epi* mask). That transformation matrix was then used to compute the GWOR_sim_ for both the best-case (*parenchymal activation*) and worst-case (*pial activation*) scenarios.

### Step 1: Results

No one registration method succeeded all the time or failed all the time; although, in general, the two cost functions that best handled the high-resolution, limited field of view data were `3dAllineate/lpc` and `FLIRT/bbr`.

The quality of alignment for each dataset was judged three ways: visual inspection by trained observers and two computations of a novel scalar metric, GWOR. One GWOR was calculated using GM_seg_, and the other was calculated using GM_act_. Each GWOR was compared against visual inspection by trained observers. Of the two, GWOR_act_ had the largest dynamic range and best agreement with observers (Fig. 3B). Alignments that received relatively low rankings from human observers had GWOR_act_ scores less than 4. In general, human rankings generally increased as the GWOR_act_ increased from 0 to 4 (Fig. 3B). For datasets with GWOR scores above 4, however, the pattern of human rankings appeared to plateau.

**Fig 3.**
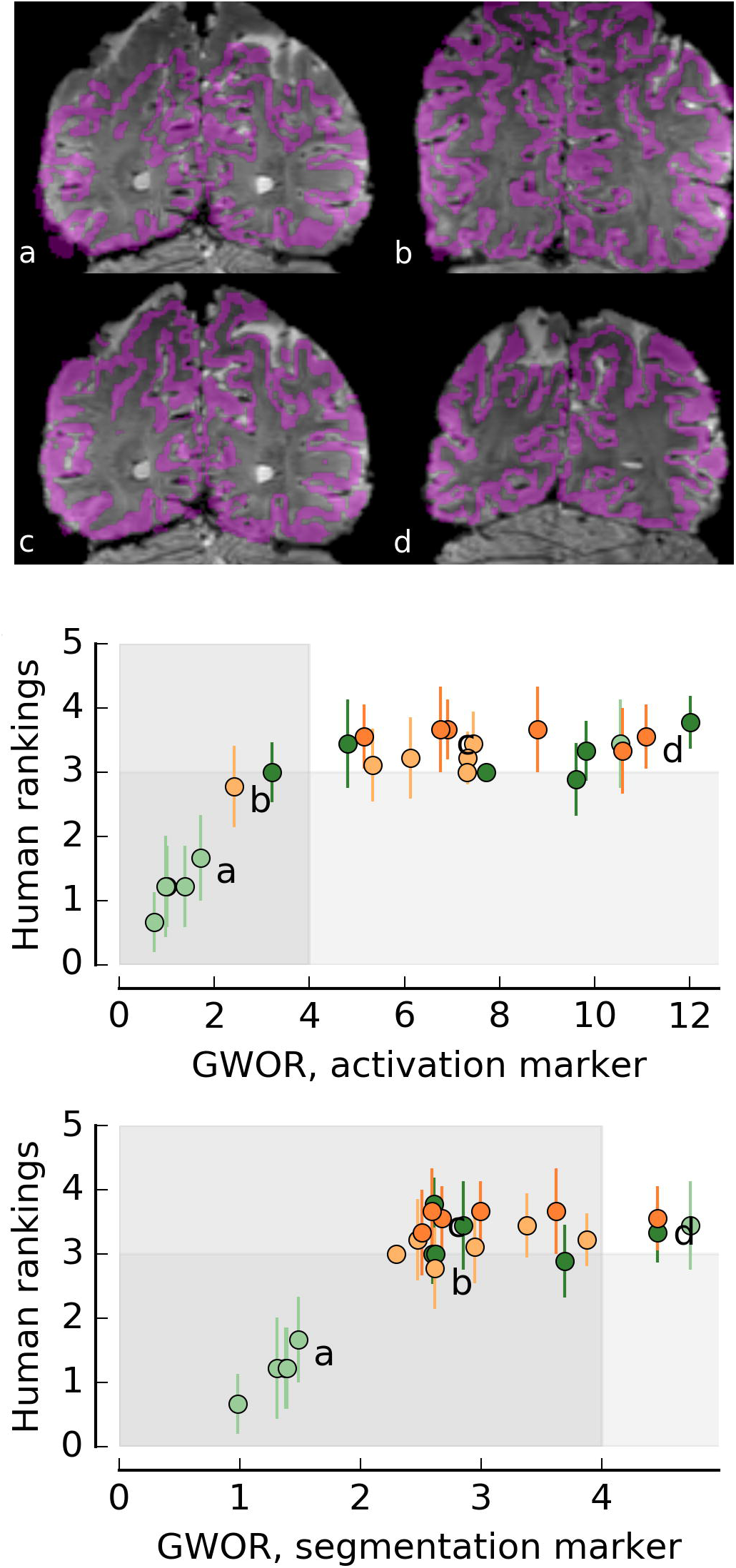
Comparison of registration quality assessed with the GM:WM overlap ratio and ranking by trained observers. (A) Representative images viewed by observers during rating. Lower case letters in (A) correspond to labeled datapoints in (B) and (C). (B) GWOR calculated for each dataset using GM_act_ compared against rankings by human observers. (C) GWOR calculated for each dataset using GM_seg_ compared against rankings by human observers. In panels (B) and (C), each datapoint represents a single registration, and different colors represent different cost functions (green = AFNI, orange = FSL). Error bars represent standard deviation.

For three datasets that contained *simulated* distortions measured in three different scanning sessions on three different individuals, the GWOR_seg_ scores were 12.6, 14.3 and 15.7. As with the real data, these numbers corresponded, upon visual inspection, to registrations with regions where the alignment appeared perfect, and small regions where the EPI images extended beyond the reference anatomy. The worst-case activation mask (*pial activation*) produced extraordinarily high GWOR_act_ scores (58-215), since it was rare that distortions were bad enough to move pial responses all the way into white matter.

### Step 1: Discussion

For the real datasets, a GWOR below 4 indicated the likely presence of observable systematic problems (as in Fig. 3B), requiring re-alignment with a different cost function, weighting function, or initial starting position before moving forward with cortex-based or surface-based analysis. Importantly, human observers were not reliably sensitive to small changes in alignment and only gave low scores when the outer boundaries of the functional and anatomical data were evidently not aligned. The particular GWOR threshold that predicts successful alignment on a given type of data will depend on the level of residual distortion in the functional data and the quality of the GM marker.

A higher GWOR score does not necessarily indicate an ideal alignment. For example, the GWOR score for the simulated pial activation dataset was very high. This instance emphasizes an important caveat for the approach: pial activation will bias scores. Thus, the GWOR is a good guide for comparing different alignments *within the same dataset*, but comparisons *between* datasets are affected by many factors, including the fact that values will be higher when activation is limited to superficial gray matter.

In any case, the GWOR is an easily computed scalar metric that largely agrees with subjective quality ratings by expert human observers, and from this first section of the work we conclude that the GWOR is useful for quantifying the quality of overall registration of functional and anatomical volumes. The GWOR can be used to automate the process of evaluating an initial registration for a given dataset, which circumvents the need for more time-consuming and subjective judgements made through visual inspection.

The Supporting Information file contains additional information about how the GWOR depends on the false positive rate, the sequestration of activation to GM, and relative volumes of GM and WM when activation is used as a GM marker. Comparison of possible scalar metrics other than GWOR against inspection by experienced observers is also shown in the Supporting Information file.

## Step 2: Region-specific masks for reliable depth-dependent analyses

In Step 1 we demonstrated a method for evaluating the overall quality of a given dataset’s cross-modal registration. Because the effects of residual uncorrected distortion are not uniform throughout a functional volume, a good *overall* alignment does not guarantee a good registration within a cortical region of interest. This problem has serious implications for performing depth-dependent analyses because alignment errors can erroneously localize superficial cortical responses to deep layers, or vice versa.

We posit that depth-dependent analysis should only be carried out in cortical areas where the *entirety* of the GM defined in the functional data overlaps with the GM in the anatomical data in which depth is defined. Restricting analyses to sections of the functional volume where functional and anatomical GM are fully aligned ensures that the experiment has the sensitivity to potentially detect activation at all depths, and, crucially, does not bias laminar profiles generated from the experimental task.

In this section, we present a pipeline for assessing the extent of the overlap between functional and anatomical GM throughout the cortical depth, from the WM boundary to the pial surface, at each surface node. This metric is called the *depth consistency fraction* (DCF). Only regions where the DCF is large (close to 1, which indicates complete overlap between functional and anatomical GM) are appropriate for further depth dependent analyses. When this method is applied to a simulated dataset, we find that laminar profiles generated from regions with poor DCF show a bias toward responses in superficial layers.

As with the GWOR score, the DCF metric can be computed using a GM marker that can be defined in different ways. In this section, we explore the use of both DCF_act_ (using GM_seg_ to define GM) and DCF_act_ (using GM_act_ to define GM). Both methods have limitations that are considered in the Discussion.

### Step 2: Material and methods

#### Computing Depth Consistency Fraction (DCF) maps

For each dataset, we used the transformation matrix resulting from the best-performing rigid-body registration (`3DAllineate/lpc`) to project each functional GM marker (GM_seg_ or GM_act_) to ten evenly spaced surfaces between the GM/WM boundary and the pial surface of the GW, using nearest neighbor resampling (AFNI’s `3dVol2Surf`). We computed the fraction of surfaces between the GM/WM boundary and the GM/pial boundary that intersected functional GM for each surface node, where all surfaces shared the same node indexing. Thus, each surface node was assigned a DCF value between 0 and 1.

#### Computing WM activation masks

It is possible that a distortion or alignment error that translates the functional data in a direction orthogonal to the banks of the sulcus will remove activation from deep layers on one side of the sulcus and introduce false activation into the WM on the other side (see the yellow circles in the bottom panels of Fig. 4). In other cases, an overlap of activation and WM is due to GM being inappropriately classified as WM (see black circles in Fig. 4). Whether the WM activation is due to an alignment flaw or a segmentation flaw, the presence of activation in the WM indicates a region where laminar analyses should not be done. Therefore, we combined the DCF metric with a mask that excluded locations in which activation extended more than 1 mm into the WM, thereby computing a WM Overlap (WMO) map. Specifically, we computed three surfaces that extended 10%, 20%, and 30% of the local GM thickness below the GM/WM boundary (on average, 1 mm into the WM). A value of 1 (i.e., overlap at all 3 meshes in the WM) was used to exclude nodes from further analysis.

**Fig 4.**
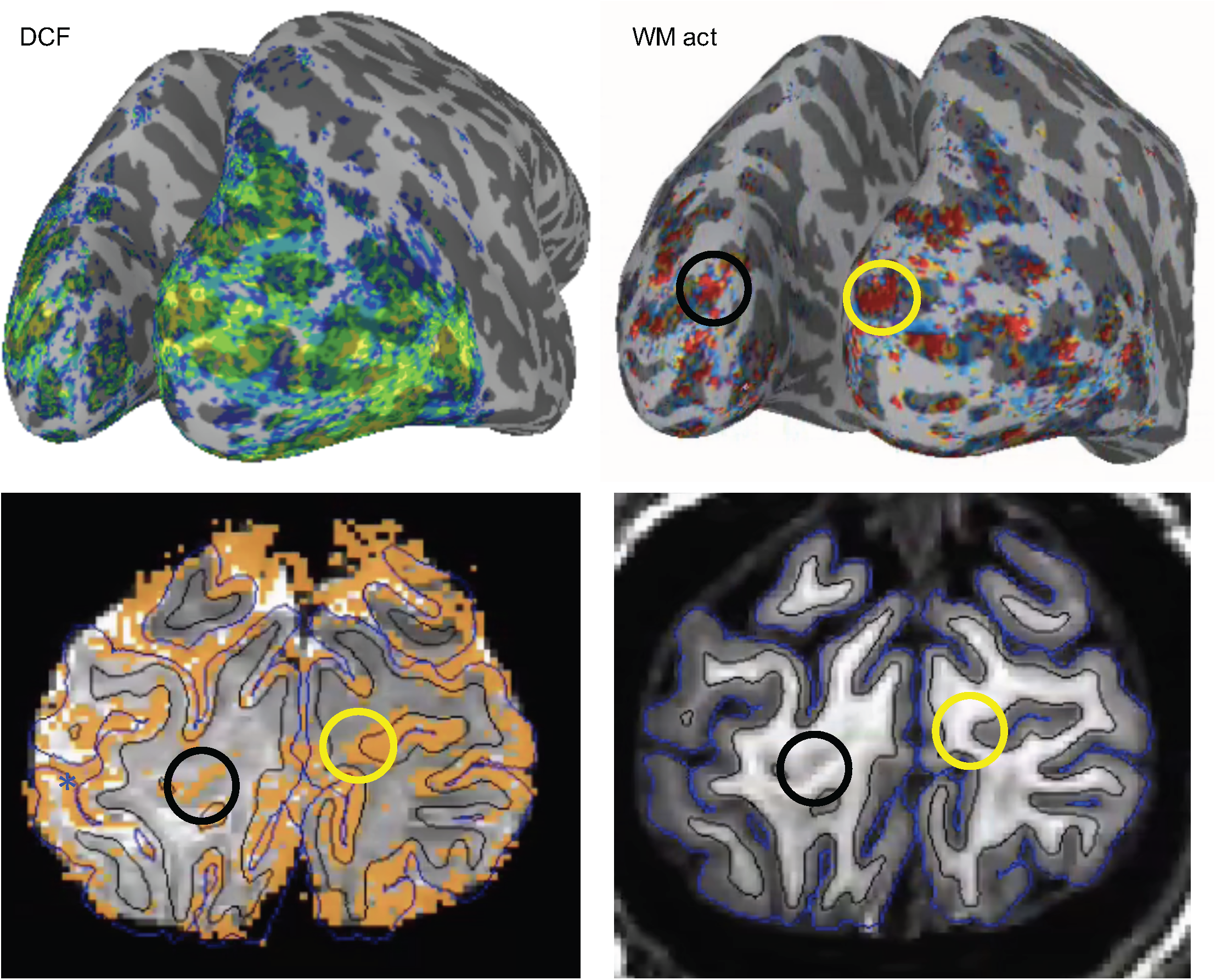
Combining GM and WM activation information to assess local alignment quality. (A) The depth consistency fraction (DCF) map indicates regions where activation is present at all depths in the GM. Poor alignment shifts activation so it is present in either superficial or deep layers. (B) The WM activation map indicates locations in which activation extended more than 1 mm into the WM. Some alignment and segmentation errors result in activation throughout the GM depth and activation in the WM, such as alignment errors that slide activation out the bottom of a sulcus (yellow circles, (C), (D)) or segmentation flaws that do not include the deepest regions of GM due to partial volume effects at the GM/WM boundary (black circles, (C), (D)). The regions of the brain representing quality local alignment are yellow or green in the DCF map and blue in the WM activation map. Subsequent laminar analyses should be pursued in these areas.

#### Computing laminar profiles

To quantify how the inclusion criteria created by the DCF maps affected subsequently estimated laminar profiles, we assessed depth-dependent responses for Experiment 2 (four block-design scans presenting high-contrast flickering checkerboards, 12 s on/12 s off per cycle, 10.5 cycles per scan). Task and independent localizer scans were co-registered so that the same transformation matrix mapped both kinds of functional data to the anatomical data (surface) space. Response amplitude estimates from a standard GLM analysis of the task scans were sampled onto a set of ten parallel surfaces spanning the GM of the Reference T_1_ using SUMA’s `3dVol2Surf`. Depth-dependent profiles were created by averaging task responses at each depth for different subsets of nodes defined by DCF computed from the independent localizer scans.

#### Simulation

The simulated data were subjected to the same process as the real data: DCFs and WMO masks were created from the ground-truth GM mask and used to score the registration quality for each node. Laminar profiles were then generated for the simulated T_2_*-weighted functional responses (ramping from 0.3% signal change at WM boundary to 1.4% at surface) as well as the flat (“parenchymal”) response profile.

### Step 2: Results

After computing DCF_seg_ and DCF_act_ maps for each of the three datasets from Experiment 2, we estimated single-condition responses for the main task and created response profiles for subsets of surface node. Ultimately, DCF_act_ and DCF_seg_ produced similar results (Fig. 5, Fig. 6). However, response amplitudes were 3-4 times larger when using DCF_act_ than DCF_seg_. The activation-based method marked a smaller volume of GM that was more likely to have robust visual responses.

**Fig 5.**
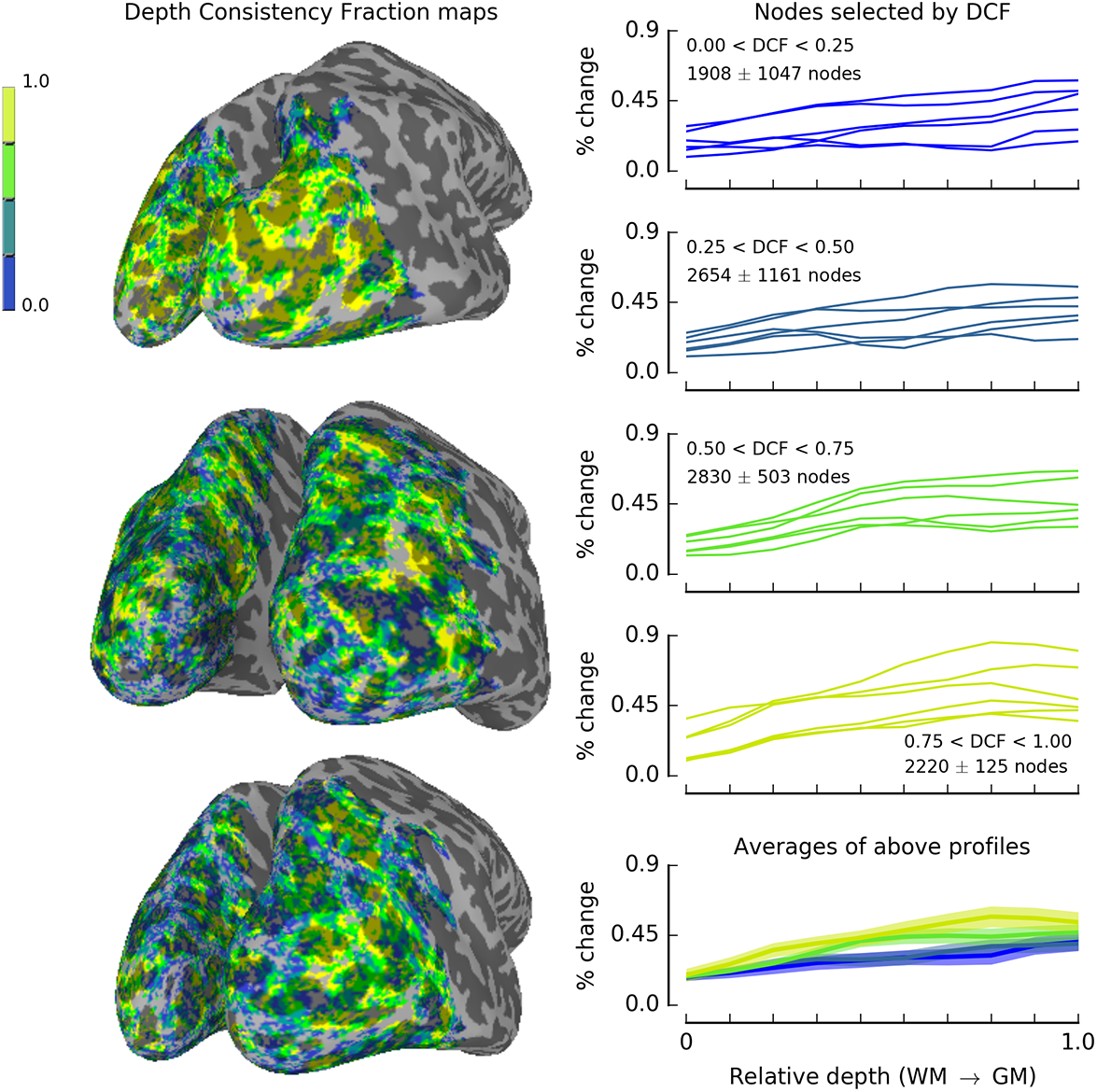
Impact of surface node selection using DCF_seg_ on depth-dependent response profiles. On the left, DCF_seg_ maps (generated by image-based segmentation in GE EPI volume) are shown for the 3 participants. On the right, laminar profiles of visual responses during the main task (checkerboard viewing) are shown for those 6 hemispheres. Top right: nodes for which DCF map showed functional GM overlapping anatomical GM in 25% or less of the cortical depth (and functional GM did not extend into the WM more than 20% of the GM depth). Fourth from top, right (yellow lines): nodes for which overlap was observed through more than 75% of the entire cortical depth. A non-overlapping set of surface nodes was used to generate each of the top 4 profiles. Bottom right: the 6-hemisphere averages for each category of surface nodes (blue: DCF < 0.25; teal: 0.25 < DCF < 0.5; green: 0.5 < DCF < 0.75; yellow: 0.75 < DCF < 1.0) are shown on the same plot for comparison, to highlight the fact that regions with poor activation/GM overlap (low DCF) create laminar profiles biased toward the pial surface. The profiles in the bottom plot do have shading indicating SEM (n=6), but the shading is comparable to the linewidth.

**Fig 6.**
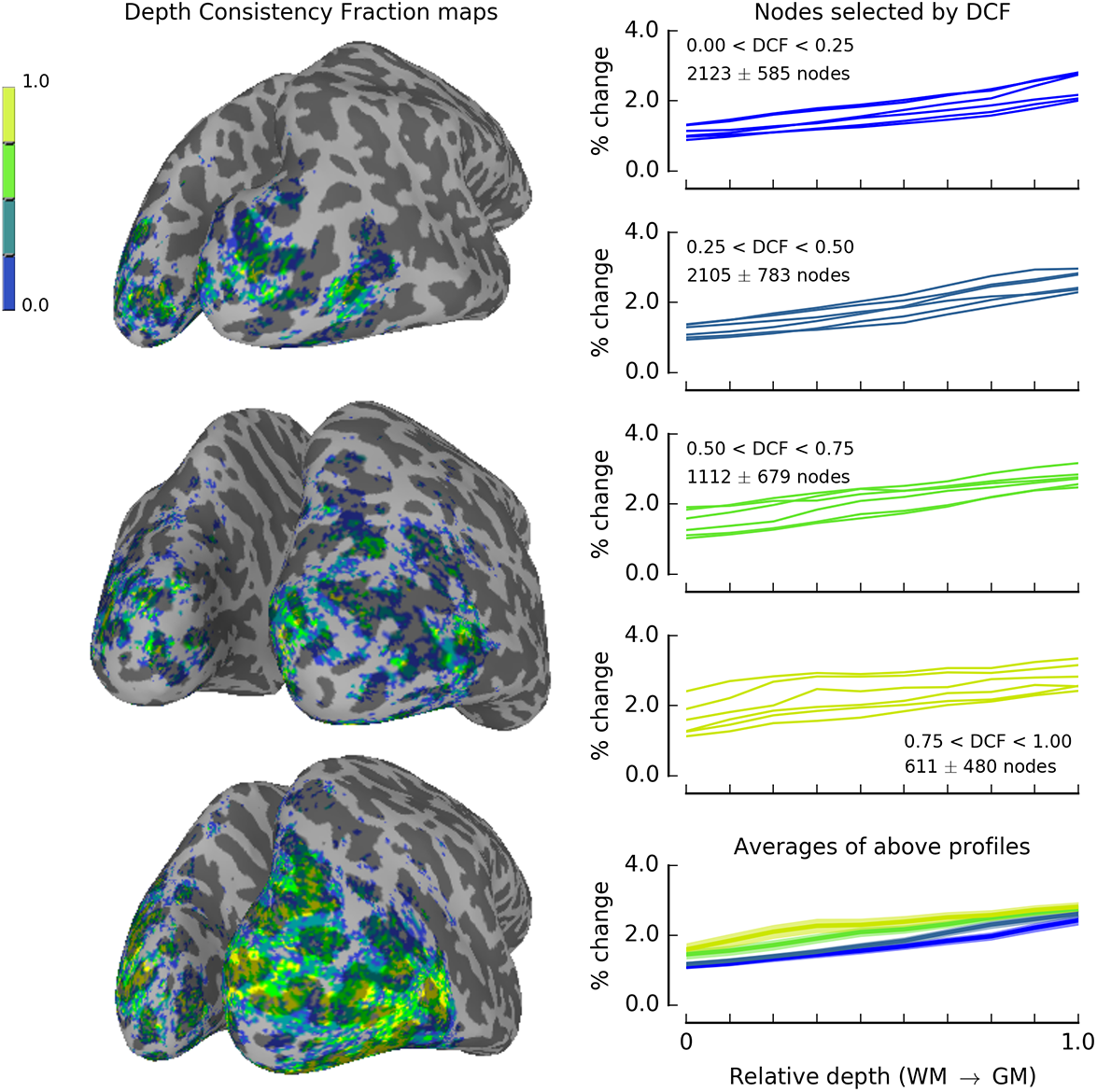
Impact of surface node selection using DCF_act_ on depth-dependent response profiles. On the left, DCF_act_ maps (generated by an independent localizer, movie watching) are shown for the 3 participants. On the right, laminar profiles of visual responses during the main task (checkerboard viewing) are shown for those 6 hemispheres. Top right: nodes for which DCF map showed functional GM overlapping anatomical GM in 25% or less of the cortical depth (and functional GM did not extend into the WM more than 20% of the GM depth). Fourth from top, right (yellow lines): nodes for which overlap was observed through more than 75% of the entire cortical depth. A non-overlapping set of surface nodes was used to generate each of the top 4 profiles. Bottom right: the 6-hemisphere averages for each category of surface nodes (blue: DCF < 0.25; teal: 0.25 < DCF < 0.5; green: 0.5 < DCF < 0.75; yellow: 0.75 < DCF < 1.0) are shown on the same plot for comparison, to highlight the fact that regions with poor activation/GM overlap (low DCF) create laminar profiles biased toward the pial surface.

Regardless of GM marker used, all of the laminar profiles reflected stronger fMRI responses near the cortex surface and reduced responses in deeper layers, as expected (see Fig. 5, 6). Importantly, using nodes for which the DCF indicated significant responses in only part of the GM depth resulted in profiles with lower responses throughout the GM thickness but relatively accentuated response near the pial surface.

The simulated data allowed us to explore this bias further. Figure 7 shows the results of the above analysis on both the flat parenchymal activation profile as well as the sloped T_2_*-weighted profile. Using the parenchymal mask to both assess alignment quality and to generate laminar profiles is of course a circular analysis that would never be done on real data, but it is illustrative in the case of this simulation. The pial bias that was observed in the real data in locations with low DCF was also observed in the simulated data, even when the simulated data was flat through the cortical depth (Fig. 7, left column).

**Fig 7.**
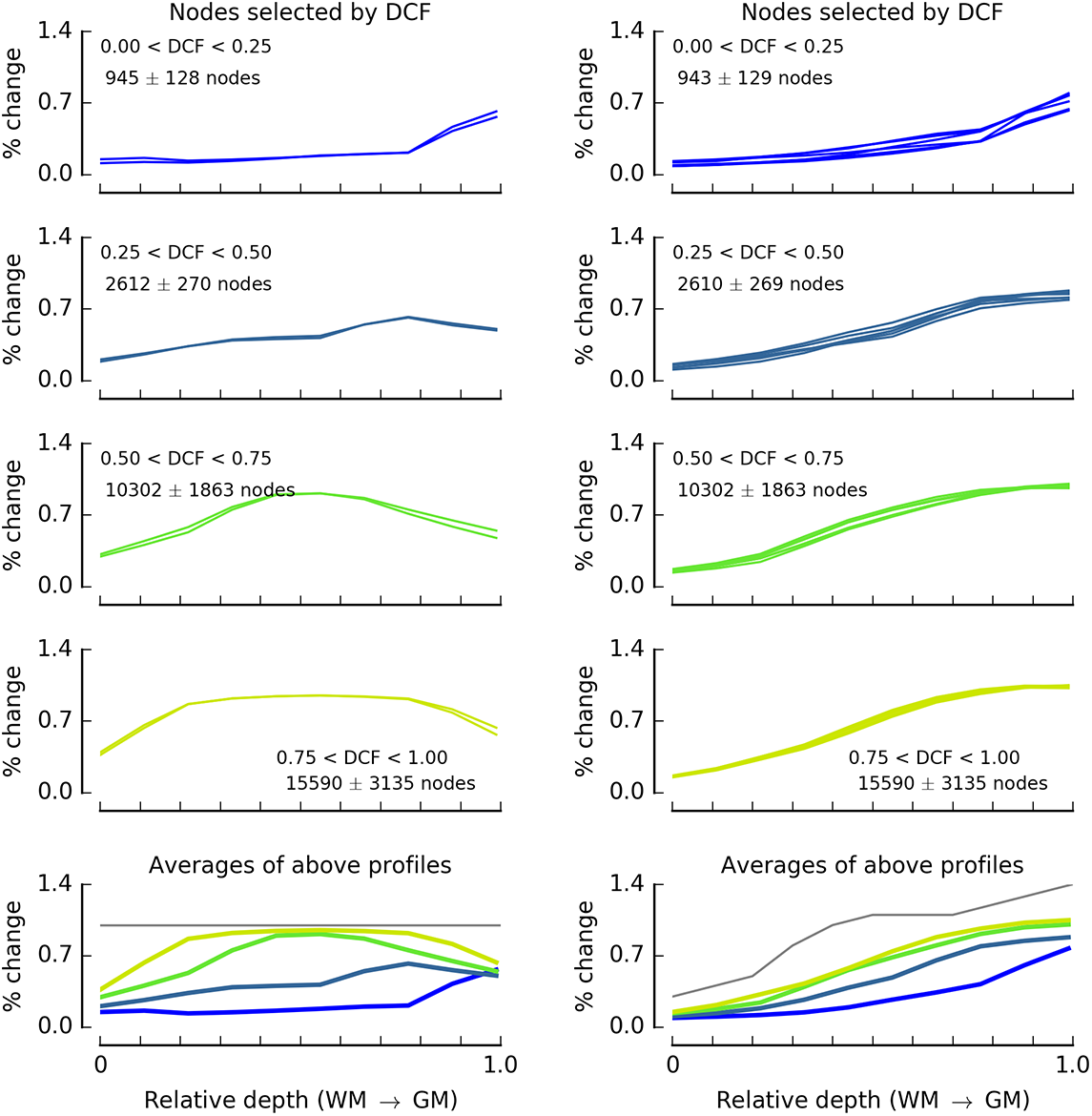
Impact of surface node selection on depth-dependent response profiles for simulated data. Left column: flat response profile throughout the GM. Right column: simulation for response profile (light gray line, bottom right plot) typical of T_2_*-weighted fMRI data. Top four plots in each column have traces from six simulated hemispheres. Colors as in Fig. 5 and 6, with light gray trace in bottom row indicating simulated profile.

### Step 2: Discussion

We explored how restricting analyses according to the *depth-consistency fraction* (DCF) would impact depth-dependent fMRI response profiles. The DCF is computed for each surface node and represents the fraction of the corresponding anatomical GM that overlaps with functional GM. Functional GM was defined with two markers: image-based segmentation (producing DCF_seg_) and activation in an independent localizer (producing DCF_act_).

The most likely explanation for the superficial bias observed in all of the generated laminar profiles is that distortion is strongest near the edges of the brain. Field perturbations could move signal into or out of the brain, depending on the sign of the phase encoding blips (R/L in our case), but we rarely observed tissue being displaced into the brain. Therefore, there was a bias for activation to appear stronger in superficial GM in places where registration was poor, emphasizing the importance of high-quality registrations in cortical regions of interest.

Although DCF_act_ and DCF_seg_ ultimately produced similar laminar profiles, the method for marking the GM has important implications. When using GM_act_ to compute DCF, it is crucial that the original task is designed thoughtfully. If the task used to compute the DCF_act_ is related to the task for which subsequent laminar profiles will be computed, the resulting laminar profiles may exhibit selection bias. Additionally, the fMRI response naturally varies in amplitude (and contrast-to-noise ratio) throughout the cortical depth. As a result, for many tasks, small but real responses are not reliably detected at all depths. Thus, when using GM_act_, it is not only important that the “localizer” task be independent from and unrelated to the task for which laminar profiles will be computed, but also that the localizer task be robust enough to generate significant signal modulation through the depth of the GM ribbon (e.g., by using a robust block-design visual localizer, motor task, or breath-hold task (ideally, the average of several scans interspersed throughout the scanning session)).

On the other hand, if GM_seg_ is used to compute the DCF, there may be some regions where segmentation errors result in too much tissue being marked as GM (i.e., WM is marked as GM). In these cases, the local DCF_seg_ value will be artificially inflated. In regions where segmentation errors are biased toward the pial surface, DCF_seg_ will be too low. Both of these errors are common in the datasets used here because the segmentation was performed on the images used for the functional studies. EPI segmentation will be better on images acquired during the same scanning session for the purpose of supporting segmentation [e.g., long TR and long TE; see 15 for excellent progress on this front]. However, even then, variability in image intensity due to tissue structure make segmentation on T_2_*-weighted EPI data challenging. Similarly, the long acquisition time required to get distortion-matched T_1_-weighted EPI images can be impractical. It is important to consider these issues when deciding how GM will be marked.

The pial bias observed in the simulated data mirror the pial bias observed in the real data, especially in locations with low DCF. This bias was evident even when the simulated data were flat through the cortical depth. It is likely that the larger effect in superficial, compared to deep, layers is related to the details of our particular datasets (coronal images in posterior cortex, with R/L phase encode). More generally, misalignment has a direct effect on the resulting depth-dependent profiles. When functional GM (and activation) overlaps partially with anatomical GM, laminar profiles will show responses only in superficial or deep layers. As a result, middle layers, on average, experience reduced responses in general. For a typical T_2_*-weighted laminar profile, with superficial responses 2-3 times larger than deep responses, the augmentation of response estimates in superficial layers will be more pronounced than in deep layers.

Another notable aspect of the laminar profiles simulated from flat parenchymal activation is the degradation of estimated response near the WM and pial surface. Here, the data end abruptly and partial-volume effects from re-sampling the data to surfaces blurs this boundary. This attenuation of response near the edges of gray matter is present in the simulated *T2* laminar profile* (the blue and green traces in the bottom right panel of Fig. 7 are relatively attenuated near the pial surface). The absence of the mid-parenchymal bump in the simulated *T2* laminar profile* reflects the resolution limits of 0.8 mm fMRI.

## General Discussion

The goal of this work was to test a two-step approach to guiding depth-dependent fMRI data analysis. The first step derives an easily computed score (GWOR) that can flag poor alignments that need more attention before moving forward in the analysis pipeline. Correspondence to ratings by expert observers verified that this was a useful screening tool. In the second step, we used the consistency of overlap between anatomical GM and functional GM throughout the cortical depth as a mask to exclude regions from further analysis. The result of this effort was the discovery that laminar profiles are biased toward superficial responses in regions where registration between functional and anatomical GM is poor, even in the case that the fMRI response is flat throughout the parenchyma.

Both steps described in this work rely on marking the GM in the functional data, which can be difficult due to low GM/WM contrast in EPI images. To address instances in which separate T_1_-weighted EPI or distortion-matched anatomical volumes cannot be acquired, we explored the use of robust activation in an independent localizer as a GM marker. Using activation as a marker for GM in the functional volume is obviously not without its limitations. If activation is more extensive in superficial layers than deep layers, then alignments in which the functional data are shifted erroneously down toward the WM would receive a higher score than alignments in which functional data were shifted erroneously away from the WM (pushing the activation up into the CSF). It is well known that fMRI sensitivity is not uniform through the cortical depth, particularly for GE EPI activation. We conclude that an activation mask with only superficial responses is an inadequate GM marker and cannot be used to assess registration quality.

A major concern that arises when using activation in T_2_*-weighted images as a GM marker (GM_act_) is that pial vessel responses, erroneously marked as GM, could bias registration. In this work, we used only rigid-body registration instead of non-linear warping between functional and anatomical data, because the GM/WM contrast in the functional data was not adequate to support non-linear registration at regions other than the high-contrast outer boundary of the brain. Thus, the tortuosity of GM protected against bias due to pial vessel activation (i.e., voxels that are actually outside of GM which are erroneously marked as GM voxels). However, if the registration algorithm allows for stretching, shearing or non-linear warping, then the inclusion of pial vessel responses in the activation-based GM marker would lead to biased registrations. Thus, we do not advocate the use of activation maps derived from T_2_* data to guide non-linear registration. Fortunately, if the image contrast is sufficient to support non-linear registration, image-based GM markers (GM_seg_) are more easily derived in the data and activation need not be used. Activation as a proxy for GM in the functional data is only valuable for datasets with poor GM/WM contrast in the EPI data and no access (either due to time constraints or scanner software) to additional distortion-matched volumes with good GM/WM contrast.

The best approach for marking GM in the functional volume is, of course, to use image contrast to segment the GM in the functional data [12,13,or distortion-matched anatomical data; for recent examples see 25]. Here, we tested a worst-case scenario of this approach in which the segmentation is performed on the functional data themselves. The fact that DCF_seg_ produced laminar profiles (Fig. 5) that agreed with the simulation (Fig. 7: reduced pial bias in regions with good DCF) indicates that even a poor EPI segmentation can support improved laminar analyses. This, as well as the fact that DCF_act_ produced laminar profiles consistent with the simulation (Fig. 6), indicates that even imperfect GM markers can guide analyses in cases where GM/WM contrast cannot be generated in the EPI data or the scanning session does not allow time to acquire an additional volume with adequate SNR for segmentation.

A natural extension of this work would be to use the functional GM marker, however it is derived, as a cost function to actually guide registration. This approach could employ existing boundary-based registration cost functions, which use image intensity change across the GM/WM boundary defined in the reference image. To test this idea, we used FSL’s `FLIRT` with a `bbr` cost function to register a volume formed from the sum of the binary *epi* mask and the GM mask to the reference anatomy. In cases where either the EPI segmentation was of high quality or the activation was very robust, the results were as good as or better than any of the others shown in this paper, thus demonstrating the feasibility of this approach. However, this approach failed in half of the datasets/GM marker combinations used in this work, indicating that only a proper GM marker derived from independent, distortion-matched scans with high GM/WM should be seriously considered for this approach.

The second half of this work addresses the problem that some regions of the functional volume are well-aligned to the anatomical reference, while others are not. This is a result of incomplete distortion compensation in the EPI acquisition. Recent studies have shown that non-linear registration between functional and anatomical data, instead of or in addition to the linear, rigid-body registration techniques used here, can produce superior results [26]. In principle, non-linear registration (such as that provided by AFNI’s `qwarp` or FSL’s `FNIRT`) could compensate for residual distortion in the EPI data that are not compensated by first-principles approaches (using field maps or PE-reversed scans to define and compensate for distortions caused by B_0_ perturbations). However, while non-linear warping is widely used and robust on a large scale (e.g., registering whole-brain data to an anatomical template), it is not as effective for fine-scale corrections, particularly on datasets with impoverished GM/WM contrast. The same mechanisms that thwart GM/WM segmentation in the functional data thwart non-linear registration algorithms. Thus, the use of a DCF map (or a DCF map in combination with a WMO map) to exclude poorly registered cortical sub-regions is a sensible solution to handling data in which residual distortions exist and cannot be corrected.

In summary, we propose that the overlap of functional and anatomical GM throughout the imaging volume, penalized by functional GM overlap with anatomical WM (GWOR), is a reasonable metric to support objective assessment of registration quality. We further promote the use of the Depth Consistency Fraction – consistent presence of functional GM markers throughout the anatomical depth – as an inclusion criterion for regions of interest when computing laminar profiles. In a recent study [27] we found this approach crucial for avoiding regions where residual susceptibility-induced distortions caused local errors in alignment. These tools, together, can support more objective assessments of data quality and interpretation of depth-dependent fMRI studies.

## Acknowledgments

We thank Emily Allen and Michael-Paul Schallmo for participating as two of our expert observers. We also thank Alexander Bratch for assistance with data collection.

## Supporting Information

**Table 1. Brief description of image acquisitions.** All data were acquired on the same scanner, but over a time that spanned several hardware changes. N: number of subjects; T_RO_: total read-out time for one image (slice). All datasets were acquired with coronal slices and repetition time (TR) of 2 s, except the third which was axial with TR=4 s. All datasets had a right ♋ ♊ left phase encode direction. Significance threshold for creating activation maps was p < 0.0001, uncorrected. Head gradient insert was AC84, (maximum strength 80 mT/m; slew rate of 33 T/m/s).

**S1 Fig. Illustration of diverse datasets used to test generalizability of GM:WM ratio and Depth Consistency Fraction for assessing alignment quality.** Top row: stimulus examples (presentation paradigm provided in Table 1). Second row: orange binary activation mask (*p* < 0.0001, uncorrected) overlaid on EPI images. Title color in each panel indicates color used to for each dataset in following figures. Third row: purple overlay indicates slice placement on representative participant. Bottom row: type of masks used to guide alignments and assess alignment quality, as in Fig. 1 of main paper.

**S2 Fig. Performance of alignment algorithms, cost functions and GM:WM ratio quality metric is similar across datasets.** See main manuscript for description of computation of GM:WM activation ratio and definition of different alignment algorithms and cost function. The inset shows that there is not a strong interaction between overall activation volume and estimated GM:WM activation ratio.

**S3 Fig. Effect of significance threshold for creating activation masks on GM:WM ratio. A**) Presuming that 10% of the GM is activated, random (IID) noise corresponding to false positive rates with *p* values from 0.000001 to 0.01 was added to activation masks simulated for a range of true GM:WM ratios in volumes with 5%-40% GM (a reasonable range for posterior coronal images). Systematic dilution of estimated GM:WM ratio was observed for thresholds of *p* < 0.0001 or higher. B) Even when a less conservative threshold is used to create binary activation masks used to compute the GM:WM activation ratio and quantify alignment quality, the relationship between expert observers’ rankings and estimated GM:WM activation ratio is unchanged.

Author Contributions
KW: Conceptualization, Investigation, Formal Analysis, Writing – Original Draft Preparation, Writing – Review & Editing
AG: Investigation, Formal Analysis, Writing – Review & Editing
PB: Formal Analysis, Writing – Review & Editing
EY: Conceptualization, Writing – Review & Editing
CO: Conceptualization, Investigation, Formal Analysis, Writing – Original Draft Preparation, Writing – Review & Editing

