## Supplementary Material for "Defining region-specific masks for reliable depth-dependent analysis of fMRI data"

**Weldon et al., Supplemental Material**

In order to test whether the utility of the proposed GM:WM ratio metric was the same for datasets with different resolution or contrast, we applied the same procedures to four other datasets: a GE EPI dataset with 1 mm (isotropic) resolution and three Spin Echo (SE) EPI datasets with 0.8 mm (isotropic) resolution. Two of the SE acquisitions were acquired with coronal orientation and one was acquired with oblique/axial slice. Table S1 details the acquisition parameters, and Fig. S1 illustrates each dataset (with the 0.8 mm GE EPI dataset from the main paper in the first row and column, respectively, for comparison).

**Table 1. Brief description of image acquisitions.** All data were acquired on the same scanner, but over a time that spanned several hardware changes. N: number of subjects; $T_{RO}$: total read-out time for one image (slice). All datasets were acquired with coronal slices and repetition time (TR) of 2 s, except the third which was axial with TR=4 s. All datasets had a right→left phase encode direction. Significance threshold for creating activation maps was $p < 0.0001$, uncorrected. Head gradient insert was AC84, (maximum strength 80 mT/m; slew rate of 33 T/m/s).

| Data description, matrix, R | N | $T_{RO}$ | RF Coil | Grad. | Localizer type | Activation threshold |
|---|---|---|---|---|---|---|
| GE EPI, 0.8 mm 200x162x56, R=3 | 6 | 54 ms | 4-ch transmit, 32-ch receive | body | Free-viewing movie (10 cyc/scan, 4 scans) | Fourier, coh > 0.35 |
| GE EPI, 1.0 mm 128x128x20, R=2 | 3 | 40 ms | 4-ch transmit, 9-ch receive | head | Mixed block design, stim vs. blank (8 scans) | GLM, F(4,1689) > 5.9 |
| SE EPI, 0.8 mm 192x144x40, R=2 | 3 | 61 ms | 4-ch transmit, 20-ch receive | head | On/off moving dots (10 cycles/scan, 2-3 scans) | Fourier, coh > 0.39 |
| SE EPI, 0.8 mm 192x144x20, R=2 | 2 | 56 ms | 4-ch transmit, 9-ch receive | head | On/off checkerboard (8 cycles/scan, 8 scans) | Fourier, coh > 0.42 |
| SE EPI, 0.8 mm 192x144x20, R=2 | 3 | 56 ms | 4-ch transmit, 20-ch receive | head | Differential localizer (10 cycles/scan, 3-8 scans) | Fourier, coh > 0.30 |

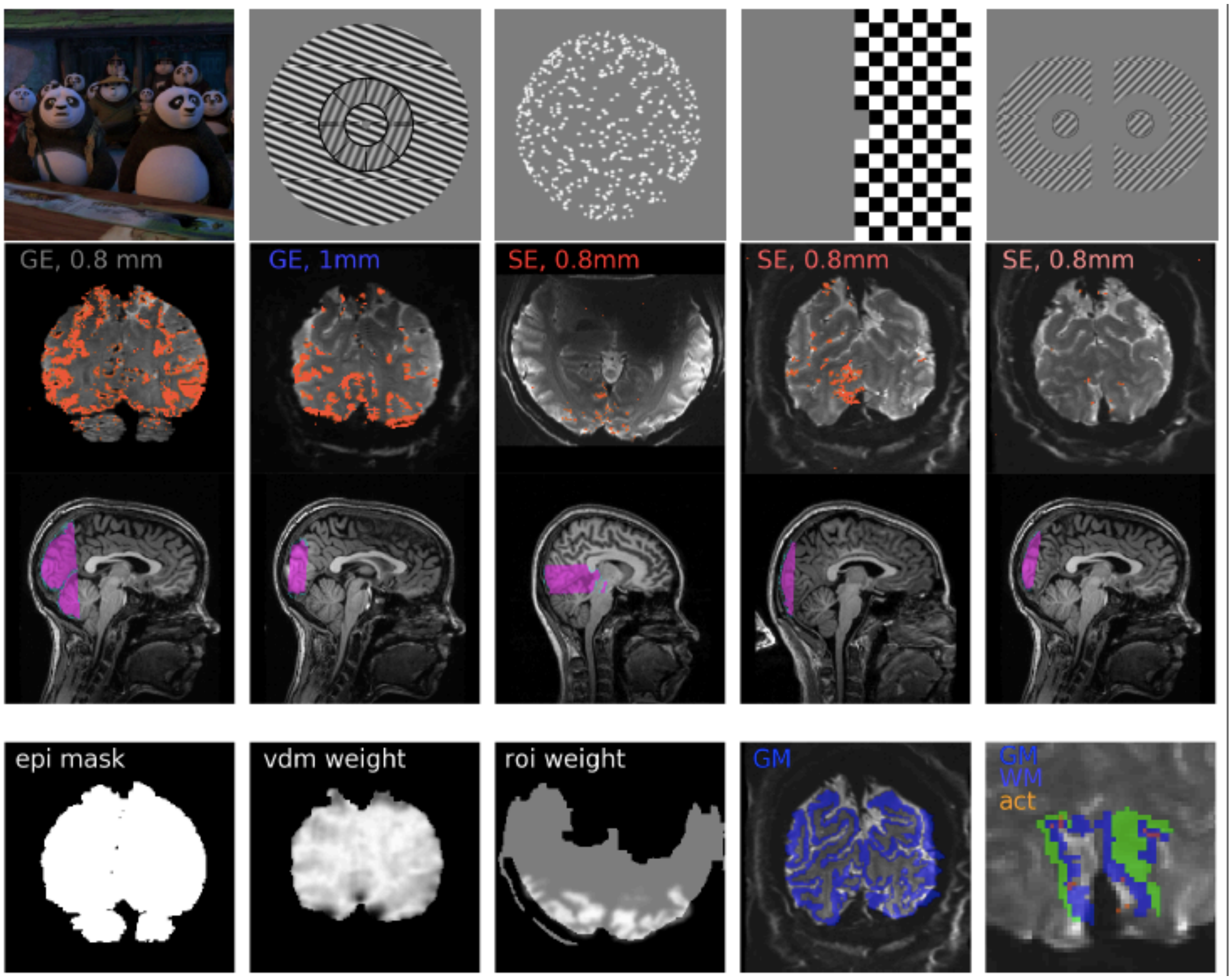


**Figure S1. Illustration of diverse datasets used to test generalizability of GM:WM ratio and Depth Consistency Fraction for assessing alignment quality.** Top row: stimulus examples (presentation paradigm provided in Table 1). Second row: orange binary activation mask ($p < 0.0001$, uncorrected) overlaid on EPI images. Title color in each panel indicates color used to for each dataset in following figures. Third row: purple overlay indicates slice placement on representative participant. Bottom row: type of masks used to guide alignments and assess alignment quality, as in Fig. 1 of main paper.

As shown in Fig. S2, performance of the different alignment metrics, as assessed by GM:WM ratio, was similar across datatypes in that FLIRT/bbr and 3dAllineate/lpc performed most reliably.

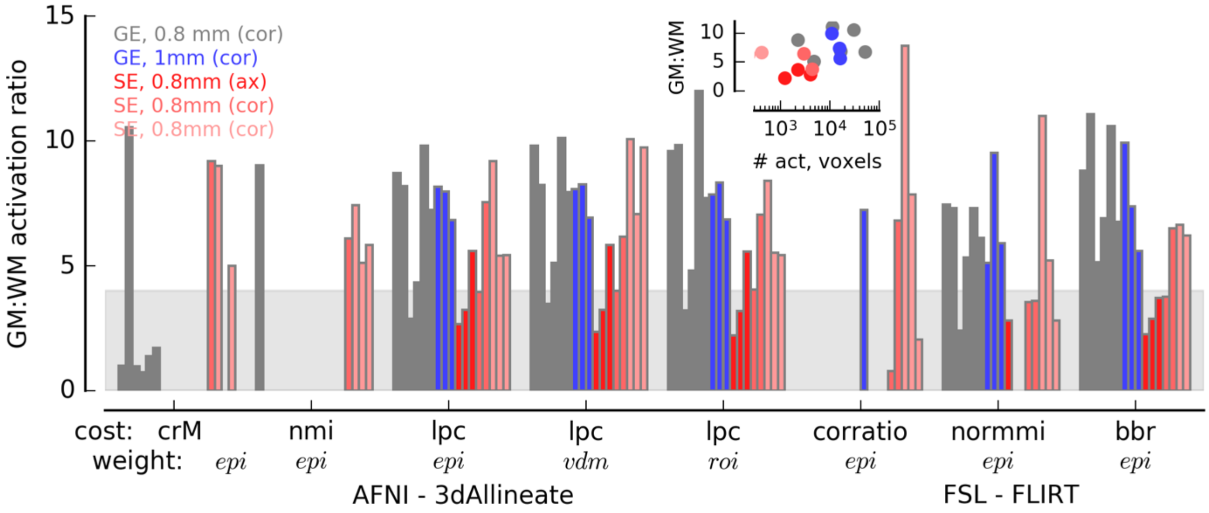


**Figure S2. Performance of alignment algorithms, cost functions and GM:WM ratio quality metric is similar across datasets.** See main manuscript for description of computation of GM:WM activation ratio and definition of different alignment algorithms and cost function. The inset shows that there is not a strong interaction between overall activation volume and estimated GM:WM activation ratio.

Because contrast-to-noise ratio is lower in SE EPI datasets than GE EPI, a threshold of $p < 0.0001$ was used to create binary activation masks for assessing alignment quality and produce the values shown in Fig. S2. This resulted in a higher rate of false positives included in the activation mask, which diluted the GM:WM ratio. Fig. S3A shows a simulation of hypothetical GM:WM activation ratios for a range of GM:WM volumes and true GM:WM ratios. This particular simulation assumes that 10% of the GM is significantly modulated; a variation of that assumption affects the GM:WM activation ratio in a way that is similar to the overall proportion of the tissue that is GM: the less GM in the image, or the less widespread the activation in the GM is, the greater the error (reduction) in estimated GM:WM activation ratio.

The logic presented in Fig. S3A also predicts that using $p < 0.0001$ instead of $p < 0.000001$ (as in the main manuscript) would reduce estimated GM:WM activation ratios for the main dataset and change the relationship between the GM:WM ratio and expert observers' ratings of alignment quality. Replotting Fig. 2B of the main manuscript in Fig. S3B (the only change is that the activation masks used to estimate GM:WM ratio were thresholded at $p < 0.0001$ instead of $p < 0.000001$) shows that the estimated GM:WM ratio is indeed reduced, especially for the highest value. But the relationship between GM:WM ratio and observers' ratings still indicates that GM:WM ratios > 4 mean that expert observers can no longer distinguish alignment quality by visual inspection.

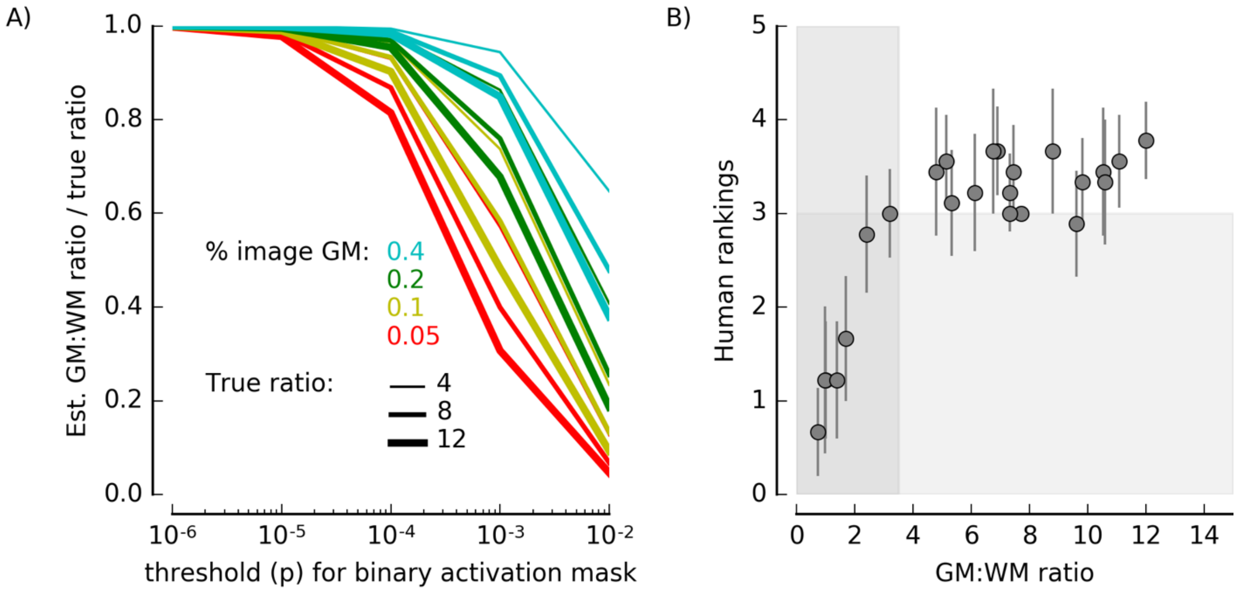


**Figure S3. Effect of significance threshold for creating activation masks on GM:WM ratio.** **A**) Presuming that 10% of the GM is activated, random (IID) noise corresponding to false positive rates with $p$ values from 0.000001 to 0.01 was added to activation masks simulated for a range of true GM:WM ratios in volumes with 5%-40% GM (a reasonable range for posterior coronal images). Systematic dilution of estimated GM:WM ratio was observed for thresholds of $p < 0.0001$ or higher. B) Even when a less conservative threshold is used to create binary activation masks used to compute the GM:WM activation ratio and quantify alignment quality, the relationship between expert observers' rankings and estimated GM:WM activation ratio is unchanged.
